# Improved motion correction of submillimetre 7T fMRI time series with boundary-based registration (BBR)

**DOI:** 10.1101/747386

**Authors:** Pei Huang, Johan D. Carlin, Richard N. Henson, Marta M. Correia

**Affiliations:** MRC-Cognition and Brain Sciences Unit, University of Cambridge, UK

## Abstract

Ultra-high field functional magnetic resonance imaging (fMRI) has allowed us to acquire images with submillimetre voxels. However, in order to interpret the data clearly, we need to accurately correct head motion and the resultant distortions. Here, we present a novel application of Boundary Based Registration (BBR) to realign functional Magnetic Resonance Imaging (fMRI) data and evaluate its effectiveness on a set of 7T submillimetre data, as well as millimetre 3T data for comparison. BBR utilizes the boundary information from high contrast present in structural data to drive registration of functional data to the structural data. In our application, we realign each functional volume individually to the structural data, effectively realigning them to each other. In addition, this realignment method removes the need for a secondary aligning of functional data to structural data for purposes such as laminar segmentation or registration to data from other scanners. We demonstrate that BBR realignment outperforms standard realignment methods across a variety of data analysis methods. Further analysis shows that this benefit is an inherent property of the BBR cost function and not due to the difference in target volume. Our results show that BBR realignment is able to accurately correct head motion in 7T data and can be utilized in preprocessing pipelines to improve the quality of 7T data.

## 1 Introduction

Participant motion is a significant confound in functional magnetic resonance imaging (fMRI) (Andre et al., 2015), and this problem is exacerbated when data is acquired at higher field strengths and submillimetre resolution (Maclaren et al., 2010). Even the best trained participants will often have unavoidable drift and unconscious motions due to respiratory (~1mm) and cardiac activity (~100um) (Maclaren et al., 2012) which can impact data quality (Hutton et al., 2011). Participant motion is a multi-faceted problem that is persistent in fMRI studies (Friston et al., 1996) and results in degrading data quality in a multitude of ways. Participant motion can affect the magnetic field, in turn causing distortions (Jezzard and Clare, 1999) and intensity variations (Friston et al., 1996) in the acquired volumes. This is a large confound since such motion artefacts affect the image in non-rigid ways and hence, standard rigid body realignment techniques might not be sufficient. Since field inhomogeneities scales with field strength, the aforementioned distortions will also become amplified at higher fields.

In addition to distortions due to field inhomogeneities, submillimetre resolution voxels also worsens the problem. While smaller voxel sizes are useful for analysis of brain substructures (e.g. cortical layers), smaller voxels also mean that any analysis would be more susceptible to motion, as the magnitude of the motion becomes larger relative to the voxel size. As most fMRI analyses assume that the same voxel corresponds to same location in the brain throughout the session. This assumption is invalidated by motion and could result in missed effects and/or false positives (Field et al., 2000; Schulz et al., 2014). Moreover, studies acquiring data at sub-millimetre resolutions generally only obtain partial brain volumes to maintain a reasonable repetition time (TR). This compounds the problem because the reduced field-of-view provides less information to drive the realignment. As such, we believe that the conventional realignment methods currently used might be insufficient to ensure the quality of ultra high-resolution data. Numerous improvements have been suggested and implemented, both at the acquisition stage (Huang et al., 2018; Todd et al., 2015) and the post-processing stage (Gallichan et al., 2016; Yarach et al., 2015).

There are two main categories of motion correction methods: Prospective Motion Correction (PMC) and Retrospective Motion Correction (RMC). In PMC, real-time motion information of the participant’s head is obtained concurrently with the acquisition of the imaging volume. This information is used to update the co-ordinates of the acquisition volume before each radiofrequency (RF) pulse to ensure that the exact same voxels are being acquired across time. Recent reviews (Maclaren et al., 2013; Zaitsev et al., 2016) provide a good overview of the PMC field and highlight some of the most promising techniques. The estimation of real-time motion information can be done by either using external tracking modules, such as an optical camera in the bore of the scanner (Callaghan et al., 2015; Stucht et al., 2015), or using the internal MR data, such as k-space navigators (Van Der Kouwe et al., 2006) or fat-based navigators (Engstrom et al., 2015). In RMC, rigid body translations and rotations are applied to each volume post-scan to align all volumes to the same scan (Ashburner and Friston, 2003). Currently, most RMC methods utilize a cost function relying on intensity differences per voxel across the volumes to drive realignment, henceforth referred to as Voxel-Based Registration (VBR) methods. There have also been attempts to address the non-rigid body nature of motion artefacts through more advanced realignment methods (Andersson et al., 2001; Chambers et al., 2015).

Both PMC and RMC come with their own set of advantages and disadvantages. PMC ensures that edge voxels are consistently captured in cases of partial brain volume acquisition and can also correct for intra-volume motion since the real-time motion information is updated after each RF pulse. However, PMC is still a relatively novel field. Specialized equipment (such as an in-bore optical camera, dentist-molded mouthpieces for marker attachment, etc) is often not widely available and implementation requires modifications to standard scanning sequences. In contrast, RMC has consistently been part of post-processing pipelines for over 15 years and has been streamlined into most post-processing pipelines, such as that in the SPM software (www.fil.ion.ucl.ac.uk/spm). It also does not require any specialized equipment. However, RMC has shown to be less than perfect, especially when compared to PMC data(Huang et al., 2018; Stucht et al., 2015).

In this paper, we propose a novel application of Boundary-Based Registration (BBR) to generate an accurate realignment of an fMRI time series to improve on conventional RMC techniques. BBR (Greve and Fischl, 2009) was originally developed to coregister images across different imaging modalities or functional contrasts, and has shown to be more effective than standard VBR methods. However, to the best of our knowledge, BBR has not been used to realign time series data. We utilized the Freesurfer (https://surfer.nmr.mgh.harvard.edu/) implementation of BBR in our realignment pipeline by coregistering each fMRI volume to the same structural volume, thereby aligning each fMRI volume to every other fMRI volume. We evaluated the performance of BBR realignment against a standard VBR approach, in this case SPM’s conventional fMRI realignment, which has been used for high-res 7T data (O’Brien et al., 2017; Tak et al., 2018).

In BBR, the grey matter boundaries are generated from a cortical surface reconstruction and are used to align the EPI image such that the maximum change of intensity is perpendicular to that of the boundary. By repeating this for each fMRI volume, this realigns the fMRI volumes to each other and to the structural at the same time. For VBR, the fMRI volumes are directly aligned to each other (without using a structural image), and importantly, the cost function is based instead on the sum of squared differences in intensity values across all voxels within a pair of fMRI volumes. We hypothesised that the fact that the BBR cost function depends only on grey-matter boundaries would benefit alignment of 7T sub-millimetre data, since it would be more robust to distortions introduced by signal inhomogeneities at medial white-matter and subcortical locations.

We conducted a visual attention task on six participants at 7T and analysed the same data using the two different realignment methods (BBR vs SPM’s VBR). We looked at four different metrics of data quality: three univariate metrics – temporal signal to noise ratio (tSNR), functional contrast to noise ratio (fCNR) and the coefficient of determination for the model fit (R2) – and the cross-validated linear discriminant contrast (LDC) as a multivariate metric (Huang et al., 2018). The stimuli were designed to probe multiple regions of interest (ROIs), in both early and higher visual areas, so as to compare the realignment methods in different parts of the brain. Furthermore, we carried out three additional realignment approaches that are intermediary between the two main methods, in order to attempt to isolate the source of any differences between the two realignment methods. These intermediary methods utilized 1) a brain mask for SPM realignment (reducing the influence of non-brain voxels on the realignment), 2) a reduced brain mask for SPM realignment and 3) realignment via coregistering each fMRI volume to the structural image, analogous to BBR, but using SPM’s between-modality, voxel-based coregistration (where the cost function is based on mutual information rather than sum-of-squares). Finally, we also applied the BBR realignment technique to 3T data from a previous study (Huang et al., 2018), in an attempt to establish whether any differences or improvements are restricted to high-field 7T data, or generalizable to other type of fMRI data.

## 2 Methods

### 2.1 7T Experiment

#### 2.1.1 7T Experimental Design

The task conformed to a 2×2 factorial design, with factors of spatial attention (along one of two directions) and stimulus category at the attended location (faces vs houses). On each trial, a participant saw two or four stimuli arranged in a 2-by-2 grid, and in different blocks, asked prompted to attend to either the positive diagonal (45° / 225°) or negative diagonal (135° / 315°). Each diagonal contained two images of the same category of stimuli, either faces or houses. Thus, there were four different types of blocks: attending to houses along the positive diagonal (H+), attending to houses along the negative diagonal (H−), attending to faces along the positive diagonal (F+) and attending to faces along the negative diagonal (F−). This allowed us to test for location selectivity and categorical selectivity by contrasting the corresponding pair of conditions, i.e. H+ and F+ versus H− and F− to investigate location selectivity, or H− and H+ against versus F− and F+ to investigate category selectivity. We expected strong location selectivity but weak or no categorical selectivity in early visual ROIs (V1, V2 and V3), and strong category selectivity but weak location selectivity in higher visual ROIs (scene-selective transverse occipital sulcus [TOS] and parahippocampal place area [PPA], plus face-selective occipital face area [OFA] and fusiform face area [FFA]).

#### 2.1.2 7T Stimuli design

All stimuli were created using Matlab (2009a, The MathWorks, Natwick, MA, USA) and presented in the scanner using Presentation (v17.2). For the main experiment, the stimuli were presented in a circular patch at four locations, diagonally from the fixation cross at 45°, 135°, 225° and 315° respectively and spanning 0.16°-2.42° eccentricity. There were a total of 20 faces and houses used as the category stimuli. All images were presented in greyscale and histogram matched across the board to equate both luminance and root mean squared contrast. This prevents any decoding due to mismatch of brightness or contrast.

A total of 20 blocks (five blocks for each of the 4 conditions) were presented during each run of the main experiment. At the start of each block, two white dots (visual angle = 0.10°) appear for 200ms indicating the pair of patches (either 45° and 225° or 135° and 315°) to which the participant should attend. This was followed by a 400ms of fixation. The stimuli then appeared for 800ms, during which the participant made a same-difference judgement between the two stimuli, followed by 400ms of fixation. This was repeated for a total of 10 trials per block, with a 1000ms rest block of fixation between each block.

In addition to the main experiment, we also acquired six runs of a population receptor field (pRF) retinotopic localizer and four runs of a categorical-selective localizer. The pRF localizer was based on (Dumoulin and Wandell, 2008; Kay et al., 2013). Both stimuli presentation and analysis scripts are available here: https://github.com/kendrickkay/knkutils/. The raw stimuli were taken from (Kriegeskorte et al. 2008). Three runs of translating bars and three runs of rotating wedges and expanding/contracting rings were presented in an alternating order. The stimulus was presented within a circular patch (radius = 7.15 °) centred on fixation with a mid-gray background.

The category-selective localizer task comprised 16-second blocked presentations of faces, scenes, objects, scrambled objects, and fixation. Each of these 5 block types appeared in a random order in each run. There was a total of 20 blocks per run (4 presentations of each block type). Within each block, 20 random stimuli from the current category were presented consecutively for 800ms each. Participants carried out a 1-back matching task while fixating on a black dot in the middle of the screen.

#### 2.1.3 Data Acquisition

Fixation is very important in this experiment to ensure that any contrast observed between blocks is not due to eye movement. Due to the inability to perform eyetracking within the 7T scanner (equipment not available), a behavioral pre-training session was carried out to ensure adequate fixation during the scan session itself, using a SMI high speed eye tracker (https://www.inition.co.uk/product/sensomotoric-instruments-iview-x-hi-speed/). Participants performed the scanner task and received feedback on their fixation performance after each run. This was repeated until the participant was able to fixate consistently (<0.05° visual angle difference between the attended and non-attended axis) for two runs.

The main experimental data was acquired on a Siemens 7 T Terra scanner using the Nova Medical 1TX / 32RX head coil. Localizer data were acquired on a Siemens 3 T Prisma-Fit scanner using a standard 32-channel head coil. Participants provided informed consent under a procedure approved by the institution’s local ethics committee (Cambridge Psychology Research Ethics Committee). A total of six healthy participants were scanned (two females, age range 25–41; two participants were authors of this study).

For the main experimental acquisition, MP2RAGE structural images were acquired first (TR = 4,300ms, TE = 1.99 ms, TI 1 = 840 ms, TI 2 = 2370ms, GRAPPA = 3, Matrix size = 320*320*224, FA 1 = 5 °, FA 2 = 6 °) This was followed by eight runs of task fMRI acquisition with the following scan parameters: 0.8mm isotropic voxels, TR=2390ms (2440ms for two participants), TE= 24ms (24.4ms for two participants), GRAPPA = 3, FA=80°, Matrix size=200*168*84, ToA=~11mins. The two participants used a longer TE and TR due to the peripheral nerve stimulation threshold being exceeded in the scanner.

For the localizer session, MPRAGE structural images were first acquired (TR = 2,250 ms, TE = 3.02 ms, TI = 900 ms, GRAPPA = 2, FOV = 256 mm*256 mm*192 mm, Matrix size = 256*256*192, FA = 9°, ToA =~ 5 min). This was followed by six runs of a pRF retinotopic localizer and four runs of a categorical-selective localizer. The EPI parameters for all localizer runs were as follows: 3mm isotropic voxels, TR=2000ms, TE=30ms, FA=78°, Matrix size=64*64*32, ToA=~5mins.

### 2.2 3T Experiment

#### 2.2.1 Experimental Design

In addition to the main 7T dataset, we investigated the effect of BBR realignment on a previous 3T dataset (Huang et al., 2018) to determine whether similar effects would be observable at other resolutions and field strengths. The 3T dataset consisted of a total of 18 participants and investigated the decoding of orientation gratings in V1.Each participant was scanned on three separate occasions (for full details, see Huang et al., 2018). The three sessions varied in whether the mouthpiece (necessary for PMC) was present (M+) or not (M−), and whether the PMC was turned on (P+) or not (P−). Note that the three sessions only consisted of P+M+, P−M+, P−M− as the fourth permutation P+M− cannot be carried out as PMC cannot be carried out without the mouthpiece. If BBR does confer any advantages in post-processing correction of motion under normal conditions, this might not be expected when PMC is turned on (P+), given that PMC has been shown to reduce motion-related artifacts (Callaghan et al., 2015; Huang et al., 2018). Within each session, the participants underwent one 11-minute run of task-based fMRI at 3mm isotropic resolution, and another run at 1.5mm isotropic resolution. The participants were asked to fixate on a blue dot in the center of the screen for the duration of the task and respond to any color changes via button press. Diagonal gratings were presented in an annulus around the fixation dot during active blocks.

#### 2.2.2 3T Data Acquisition

The 3T data were acquired on a Siemens 3 T Prisma-Fit scanner using a standard 32-channel head coil. Imaging parameters for the 3mm isotropic EPI were: TR=1260ms, TE=30ms, FA=78°, Matrix size=64*64*20, ToA=~11mins. Imaging parameters for the 1.5mm isotropic EPI were: TR=3050ms, TE=30ms, GRAPPA=2, FA=78°, Matrix size=128*128*40, ToA=~11mins. Field-of-view (FOV) parameters for both 3mm and 1.5mm EPI sequences were chosen such that the same volume was imaged across scans. For further details on the 3T study, refer to (Huang et al., 2018).

### 2.3 Data Analysis

The data first set underwent temporal interpolation in SPM 12 to correct for differences in slice acquisition times. The images then underwent distortion correction using TOPUP in FSL. The distortion was calculated using five reverse phase encode (PE) images, acquired at the start of each run, and the first five images of the fMRI time series. Distortion correction was not carried out for the 3T images.

#### 2.3.1 Realignment Methods

After initial pre-processing, the volumes then underwent five different realignment methods: two main methods and three subsidiary methods.

##### Main Realignment Methods

The two main methods were functional-structural BBR realignment and functional-functional VBR realignment in SPM. For functional-structural BBR realignment, we applied the Freesurfer implementation of the BBR function in a two-step process. First, the fMRI images were averaged across volumes, and the mean fMRI image aligned to the structural using BBR to generate an initial realignment matrix. Next, each fMRI volume was aligned to the structural using BBR with the initial realignment matrix as the seed, to reduce computation time and to reduce the probability of convergence failures due to local minima. This operation combines motion correction of functional images with coregistration to the structural image in one processing step.

For functional-functional VBR realignment, we used standard rigid-body realignment based on a sum-of-squares cost function as implemented in SPM12, which we call here the functional-functional VBR approach. Previous studies (Morgan et al., 2007; Oakes et al., 2005) have shown that while there are subtle differences between the various software packages (SPM, Analysis of Functional Neuroimages (AFNI), BrainVoyager and FMRIB Software Library (FSL)), the packages all perform similarly overall. To minimize resampling of the functional data, the structural was then coregistered to the functional data using BBR, analogous to the BBR fMRI method above. The same transformation was applied on the ROIs mentioned in Section 2.3.2 to map them into functional space.

##### Subsidiary Realignment Analyses

To further probe the cause of the differences between the two main realignment methods, we evaluated three additional realignment methods. First, we performed a variant of the functional-functional VBR realignment method where the motion estimation was restricted to a full brain mask (both shaded areas in Figure 1). Second, we carried out a similar analysis with a smaller brain mask (the red area in Figure 1), in which the cortical surface was further eroded (functional-functional VBR realignment with small brain mask). The definition of both masks is described in Section 2.3.3. Lastly, we repeated the BBR realignment pipeline, however, utilizing SPM between-modality coregistration instead of the BBR coregistration. We refer to this realignment method as functional-structural VBR. In this method, we realign every fMRI volume at each timepoint to the structural using the default normalized mutual information cost function.

**Figure 1:**
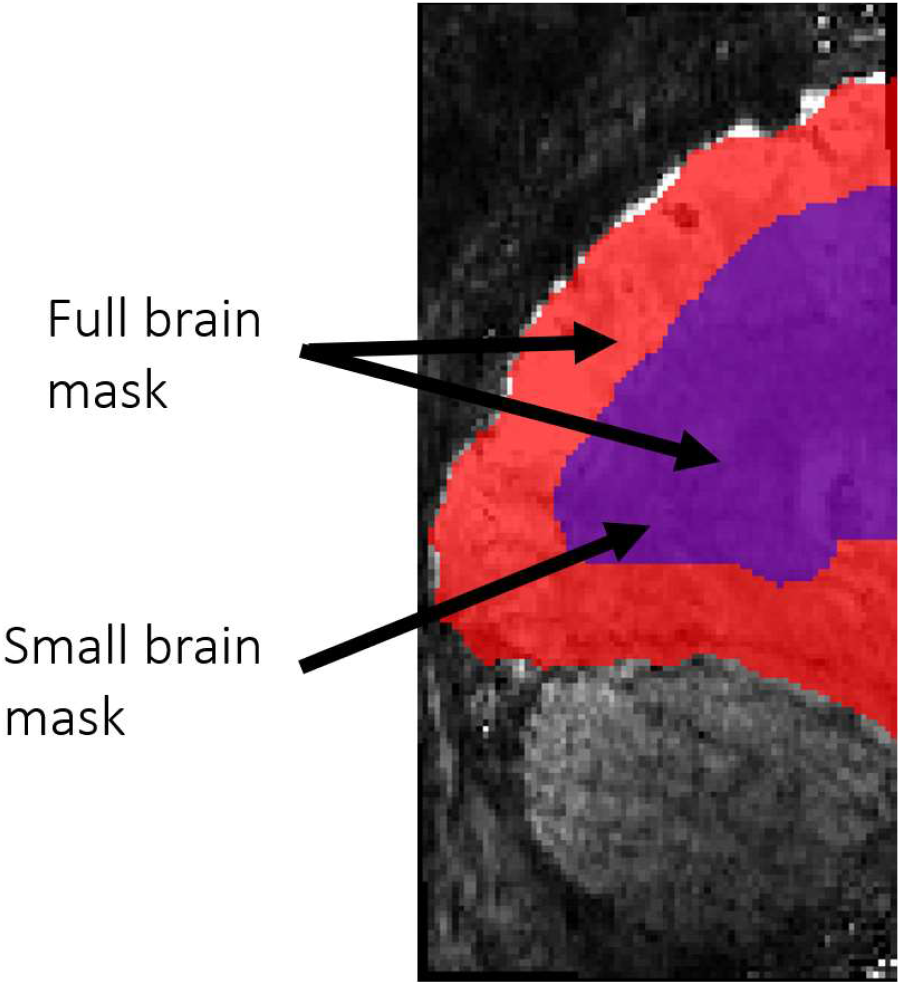
An illustration of the two brain masks utilized for the subsidary realignment methods. The full brain mask consists of both the red and purple areas while the small brain mask consists of only the purple area.

The two functional-functional VBR realignments with brain masks serve to remove the potential confound of non-brain voxels harming the standard SPM realignment. Note that since BBR realignment is driven solely by boundaries, this method already ignores out of brain volumes. A smaller brain mask (in which more voxels outside the brain surface were removed; see below for details) was also utilized because anecdotal reports have shown this method to result in better realignment. Comparisons with the functional-structural VBR realignment isolates whether differences between the two main methods are due to a methodological difference (realigning within a time series vs realigning via a structural template) or whether the benefit is inherent to the different cost functions (brain features) used.

We hypothesize that masking out-of-brain voxels should improve the accuracy of the realignment relative to standard functional-functional VBR realignment without masking. In contrast, we expect that function-structural VBR to perform worse than that of any other realignment methods. This is because SPM’s realignment with a sum-of-squares cost function (utilized in functional-functional VBR) is specifically developed for the purposes of realigning within a functional time series, and so should be superior to SPM’s coregistration with a Normalized Mutual Information cost function (utilized in structural-functional VBR), which is designed for cross-modality realignment.

#### 2.3.2 Regions of interest (ROIs)

Seven distinct ROIs were analysed: three retinotopic (V1, V2, V3) and four category-selective (FFA, OFA, pFs and TOS).

Retinotopic ROIs were defined in Freesurfer 6.0.0. Retinotopic activation maps were generated from the retinotopic pRF localizer. The maps were then projected onto a polygon-mesh reconstruction of the individual participants’ cortical surfaces. These maps were then used to manually segment out V1 to V3 on the cortex. Each individual visual area was also manually segmented into ventral and dorsal regions (e.g. V1 into V1v and V1d) for the purposes of fCNR analysis.

For the category-specific ROIs, activation t-maps where obtained using SPM by fitting a GLM to the fMRI data from the categorical localizer runs. The face-selective areas (FFA and OFA) were obtained from a contrast map by subtracting the object activation t-maps from the face activation t-maps. Similarly, the scene-selective areas (TOS and PPA) were obtained from a contrast map by subtracting the object activation t-maps from the scene activation t-maps. For each ROI, we took the 100 most activated contiguous voxels in regions that correspond to their expected locations on a brain atlas.

As all localizer data were obtained at 3T, coregistration to the 7T dataset was needed. This was done by first coregistering the 3T functional data to the 3T structural data using SPM’s between-modality coregistration. The 3T structural data were then coregistered to the 7T structural data, again using the SPM coregistration. The transformations from both coregistration steps were then applied to the ROI data. For the BBR realignment method, no further transformation was necessary since the BBR realignment process realigns the functional 7T data to the structural data. However, for the standard SPM realignment method, BBR was used to coregister the 7T structural data to the 7T functional data and the corresponding transform was applied to the ROI data. All ROI data were only resliced once all transforms have been applied.

#### 2.3.3 Brain Masks

The full brain mask was obtained by combining the grey matter and white matter voxels from the Freesurfer reconstruction and was coregistered to the functional volumes using BBR. Then the full brain mask underwent dilation and erosion by two voxels to fill in the sulci voxels. The small brain mask was obtained by eroding the full brain mask by 10 voxels. A sample volume of both brain masks superimposed on the mean fMRI image is shown in Figure 1.

We also created a grey matter mask solely for tSNR analysis. This was obtained by combining all the grey matter voxels from the Freesurfer reconstruction and coregistering it to the functional volume using BBR.

### 2.4 Data Analysis

The processed data were analyzed using the following four metrics (tSNR, fCNR, R2 and LDC):

#### 2.4.1 tSNR

The tSNR for each voxel was obtained by dividing the mean voxel intensity across the entire time course by the standard deviation of the voxel intensity. The mean tSNR was obtained by averaging across all the voxels in the grey matter mask. In the further analysis, the mean tSNR of the central brain region was obtained by averaging across all voxels in the small brain mask.

#### 2.4.2 fCNR

The postprocessed fMRI data was fit by a GLM that modeled responses to each of the four different attention conditions. For each block in the GLM, a boxcar model was used and then convolved with the canonical SPM HRF. Motion covariates were not included in the GLM. Linear and first-order sinusoidal detrending were applied to the data to remove signal drift. For each voxel, the fCNR was calculated by dividing the contrast of interest by the standard deviation of the noise. For the early retinotopic ROIs, the contrast of interest was obtained by subtracting the two conditions where the stimulus was absent in the retinotopic area from the two conditions where the stimulus is present (i.e. H+ and F+ against H− and F− or vice versa depending on the ROI). For categorical ROIs, the contrast was obtained by subtracting the two conditions where the stimulus was of the other category from the two conditions where the stimulus was of the category for which the ROI was selective for (i.e. H+ and H− against F+ and F− or vice versa depending on the ROI). The standard deviation of the noise was obtained from the standard deviation of the residuals of the GLM.

#### 2.4.3 R2

The R2 value was calculated from same GLM as for the fCNR analysis. For each voxel, the R2 was obtained by dividing the variance of the model fit by the total variance of the processed data. The model fit was obtained by multiplying the beta estimates for the four attention conditions with the design matrix. This represents the percentage of variance explained by the model. A higher R2 value indicates that the model is able to explain more of the variance in the data.

#### 2.4.4 LDC

In addition to the three univariate measures, we also investigated the effect of the different realignment methods on multivariate activation patterns using the cross-validated LDC. The cross-validated LDC (Kriegeskorte et al., 2007; Walther et al., 2016) is a contrast estimate between two conditions measured using a discriminant, which is made up of a weighted combination of the ROI voxels. An independent set of data was used to generate the weights so as to maximize the sensitivity of the LDC to differences between the two conditions of interest. Cross-validation was performed to remove the positive bias in the distance estimate due to noise (which is by definition positive) (Walther et al., 2016). This measure is also referred to as the cross-validated Mahalanobis (crossnobis) distance (Kriegeskorte and Diedrichsen, 2016).

For this paper, all presentations of each condition (H+, F+, H−, F−) from three of the four task runs were modeled as a single regressor in the design matrix. Both the data and the design matrix underwent linear and first order sinusoidal detrending. The detrended data matrix was then fit to the detrended design matrix to generate contrast estimates for the four block types. For the early retinotopic areas, we contrasted the H+ and F+ blocks against the H− and F− blocks to produce the representational distance metric. For the categorically selective areas, we contrasted the H+ and H− blocks against the F+ and F− blocks to generate the representational distance metric. This representational distance metric was normalized using the sparse covariance matrix of the noise residuals to produce the weights vector from the independent data set (Ledoit and Wolf, 2003). The data from the remaining task run underwent the same detrending and fitting to generate a test contrast estimate. The LDC test statistic is calculated by taking the dot product of the representational distance metric and the test contrast estimate. We repeated this procedure four times, utilizing a different task run to generate the test contrast estimate for each iteration. We averaged across the four LDC test statistics to generate a final continuous performance estimate, which is centered on zero under the null hypothesis of no reliable differences between the two groups of conditions. As the number of voxels used in this analysis varied across participants and ROIs, the LDC was normalized by dividing the metric by the square root of the number of voxels.

#### 2.4.5 Wilcoxon signed-rank test

Since six datapoints (participants) is not sufficient to check the normality assumption of Gaussian error that is assumed by parametric tests, the tSNR, fCNR, R2 and LDC data were analysed using a non-parametric, pairwise Wilcoxon signed-rank test. The Wilcoxon signed-rank test only requires that the data is on an interval scale and each pair of observations are random samples from a symmetric distribution. Significance was defined with an alpha level of .05. Due to the small sample size (six participants), our results will only be significant (p=0.0313) if all six participants demonstrate changes in the same direction. In all other cases, the results would not be significant (p>0.0625).

## 3 Results

### 3.1 Analysis of the two main realignment methods

#### 3.1.1 tSNR analysis of 7T fMRI data

We analyzed the tSNR for the two main realignment methods (see Figure 2, Panel A for a comparison map between the two methods on a sample participant). Improvements due to BBR realignment were heavily localized on the brain surface, which concurs with our expectations since BBR is boundary driven. In central regions of the brain, there is no visually discernable advantage of any methods, and the voxels showing preference for either method are most likely reflecting random fluctuations.

**Figure 2:**
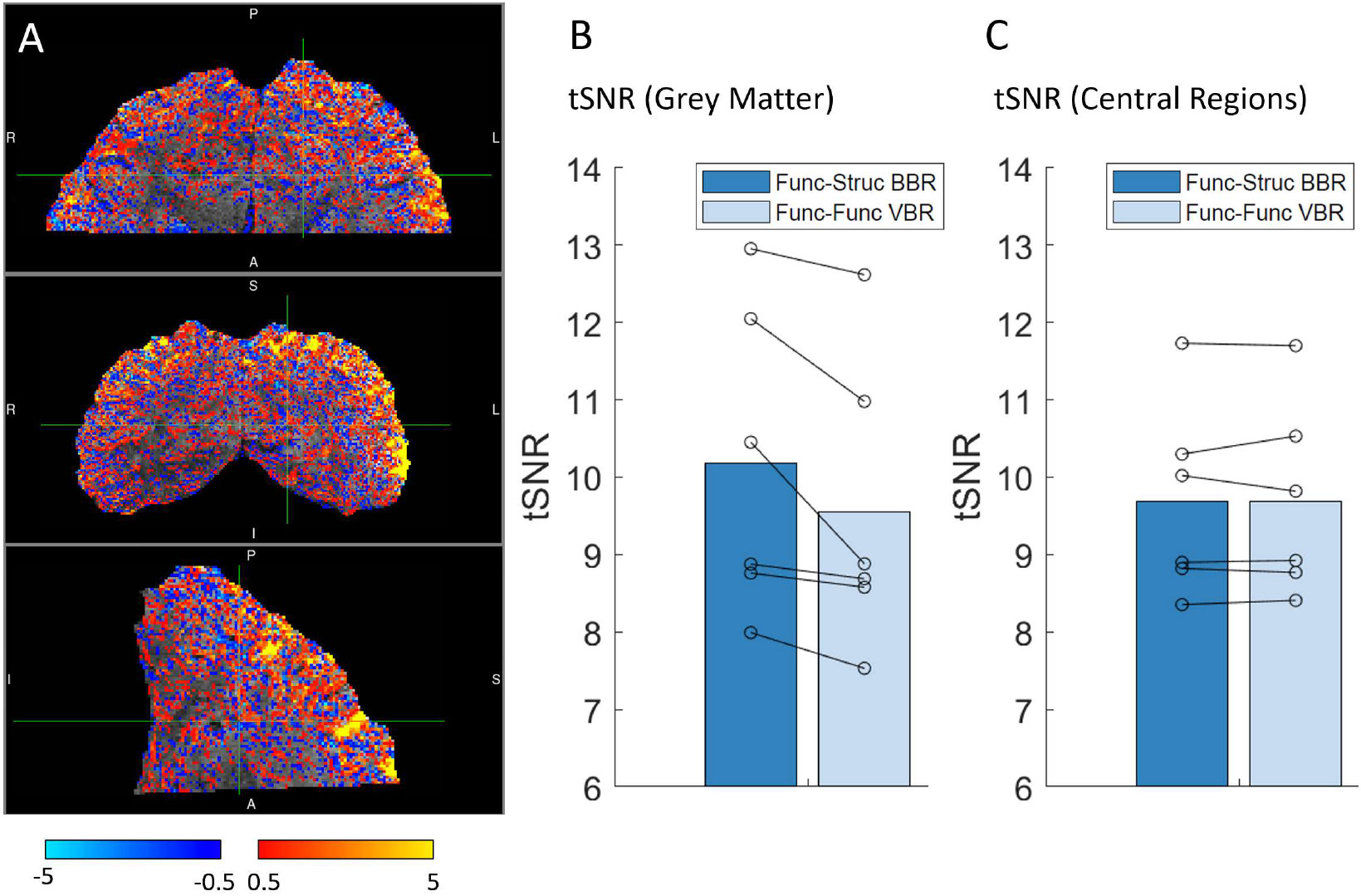
Panel A shows a comparison of the tSNR of the two main methods using a sample participant. The heatmap is generated by subtracting the functional-functional VBR tSNR from the functional-structural BBR tSNR. Red-yellow areas indicate regions where functional-structural BBR performs better while blue-teal areas show regions where functional-functional VBR performs better. Panel B shows the tSNR of the grey matter across all 6 participants when the two main realignment methods were used. Panel C shows the tSNR of the two main realignment methods when the small brain mask (Figure 1) is applied. For both Panel B and C, each pair of connected circles indicate single participant results while the bar shows the group average.

By averaging over all grey-matter voxels, we found that BBR significantly outperforms VBR under the Wilcoxon signed-rank test (Figure 2, Panel B). In contrast, averaging over the central (mostly white matter) brain regions using the small brain mask, both methods yielded very similar results (mean=9.69 for both, Figure 2, Panel C). These results are consistent with what we observed from the heatmap in Figure 2A and with our expectations that BBR realignment are more beneficial towards voxels near the boundaries.

#### 3.1.2 fCNR analysis of 7T fMRI data

Analysis of fCNR in visual ROIs provided further evidence that BBR realignment outperforms the standard VBR approach (Figure 3, Panel A). When the fCNR was averaged across all ROIs within each participant, the Wilcoxon signed-rank test indicated that BBR realignment is significantly benefits our data relative to VBR realignment. For individual ROIs, only V1 and V2 ROIs showed significant differences under Wilcoxon signed-rank testing. All other ROIs demonstrated a general trend of BBR realignment being better than standard VBR realignment, although this improvement is not consistent across all participants.

**Figure 3:**
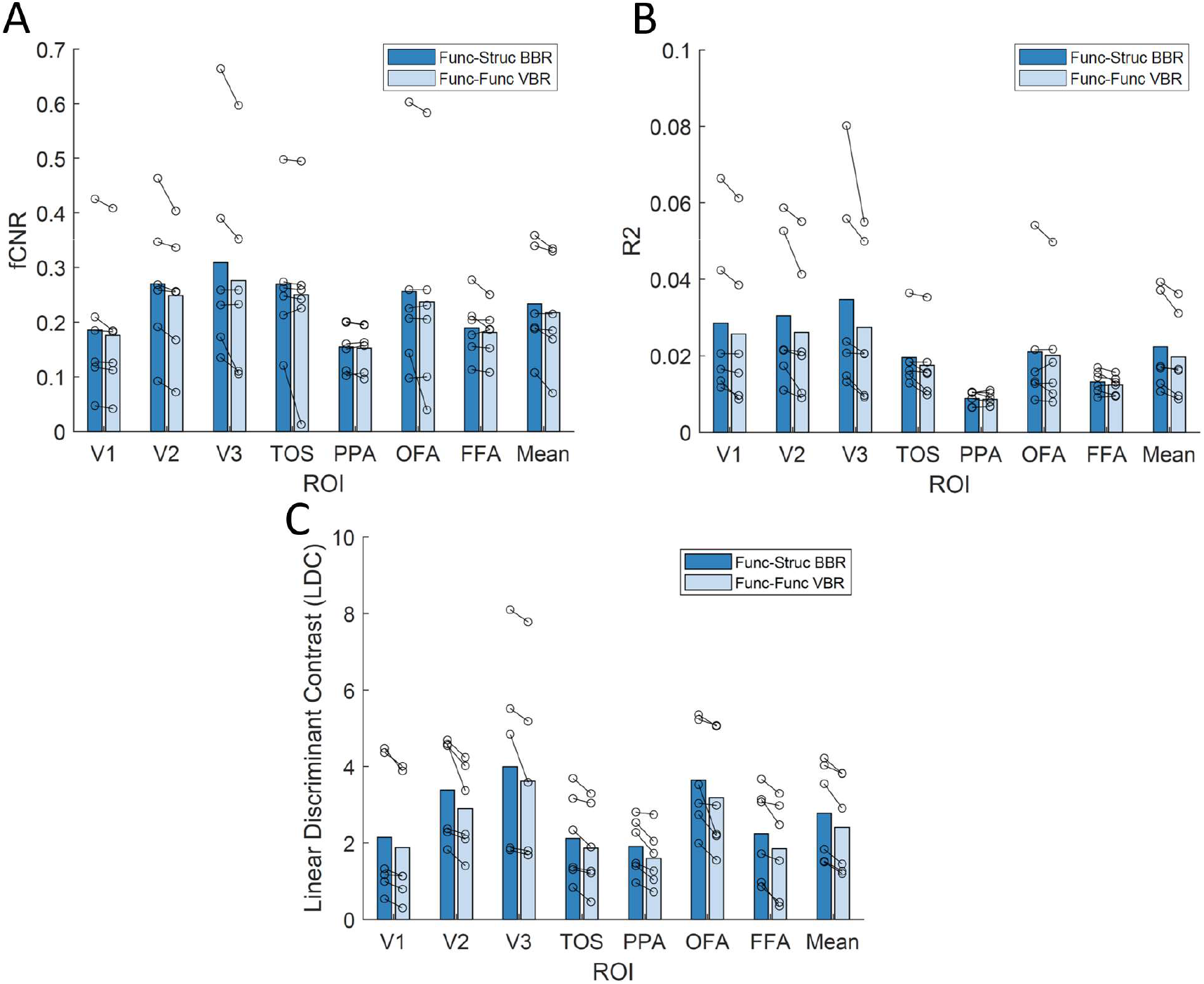
These plots compare functional-structural BBR against functional-functional VBR across multiple metrics— fCNR (Panel A), R2 (Panel B) and Linear Discriminant Contrast (Panel C). Each pair of connected circles indicate single participant results while the bar shows the group average.

#### 3.1.3 R2 analysis of 7T fMRI data

The R2 results are shown in Figure 3, Panel B and exhibit a very similar trend to that of the fCNR results, as is expected since these metrics are closely related. Averaging the R2 results across all ROIs within each participant shows that BBR realignment significantly outperforms VBR realignment under the Wilcoxon signed-rank test. Individual ROI results show significant differences for V1, V3 and TOS while all other ROIs show a small, but non-significant, benefit of BBR realignment over standard realignment.

#### 3.1.4 LDC analysis of 7T fMRI data

The LDC results are plotted in Figure 3, Panel C. Similar to the R2 and fCNR results, the average LDC across all ROIs showed a significant improvement under Wilcoxon signed-rank test when BBR realignment was used. Moreover, for all individual ROIs, the LDC from the BBR realignment data was significantly higher than that of VBR realignment data. This suggests that there is a consistent benefit of BBR realignment across all ROIs.

#### 3.1.5 tSNR analysis of 3T fMRI data

Applying the two main realignment methods to the 3T data tells a very different story (Figure 4). At 1.5mm (Figure 4, Panel A), both methods showed very similar tSNR results across all three sessions and no significant differences were observed when the Wilcoxon signed rank test was carried out (p=0.47). At 3mm (Figure 4, Panel B), BBR realignment performed significantly worse than standard VBR realignment for all 3 sessions (p = 0.00049 under Wilcoxon signed rank test). Note that these findings applied regardless of whether PMC was active (P+) or not (P−), and whether a mouthpiece was present (M+) or not (M−); see Huang et al. (2018) for details of 3 conditions. These results indicate that there is no benefit to using BBR for 3T data and it could even be detrimental (in the case of 3mm isotropic fMRI data). Given that we did not observe any benefit at the level of tSNR, we did not carry out further analysis with the other metrics or subsidiary methods. Moreover, due to the differences in the nature of the task, we would be unable to make any meaningful comparison between the 3T and the 7T data for fCNR, R2 and LDC.

**Figure 4:**
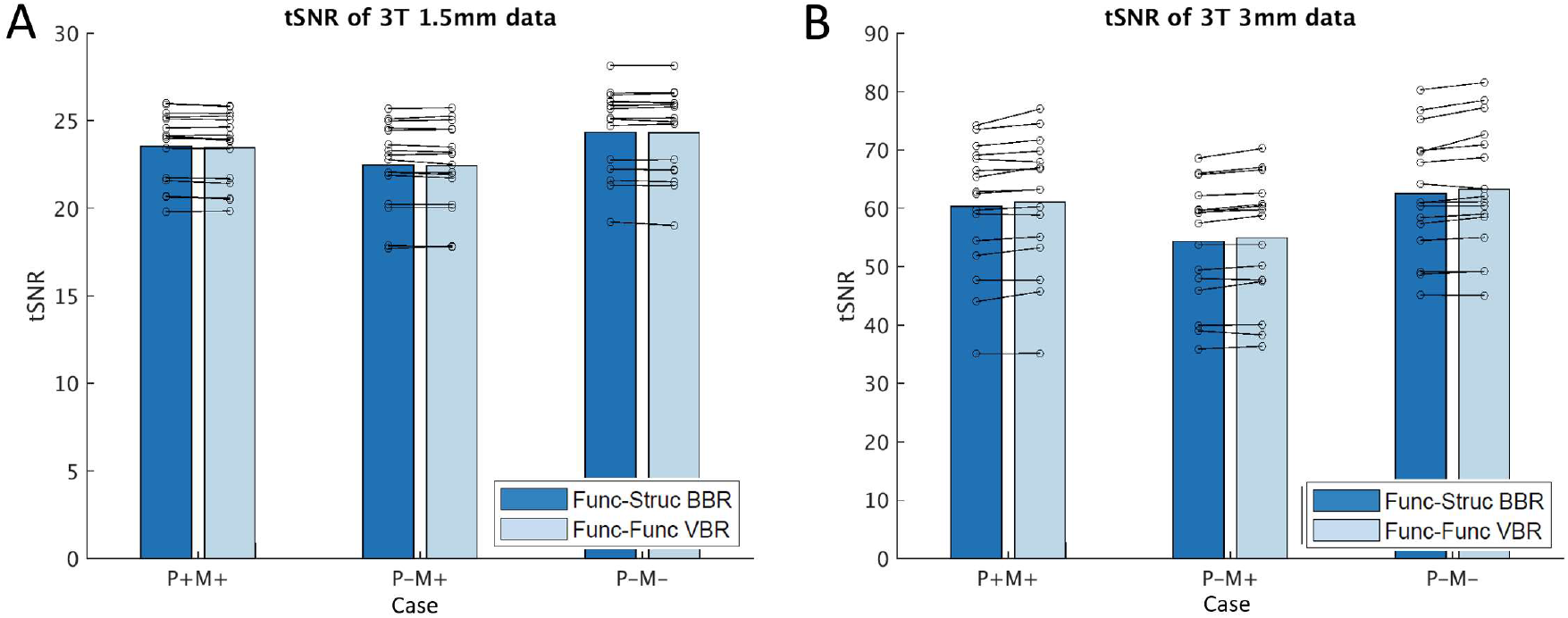
These plots compare the tSNR of functional-structural BBR against functional-functional VBR for 3T data at 1.5mm isotropic resolution (Panel A) and 3mm isotropic resolution (Panel B). The three cases on the x-axis corresponds to the type of PMC used- PMC On, Mouthpiece On (P+M+); PMC Off, Mouthpiece On (P-M+); PMC Off, Mouthpiece Off(P-M−). The fourth condition (PMC On, Mouthpiece Off) was not carried out because PMC was unreliable in the absence of the mouthpiece. Each pair of connected circles indicate single participant results while the bar shows the group average.

### 3.2 Subsidiary analyses

To probe for the source of the difference between the VBR and BBR realignment results, we designed three subsidiary analyses to help bridge the gap between the two main analyses. These analyses are not included in Section 3.1 as they are not standalone methods of improving fMRI realignment, but rather a way of understanding the differences between the functional-functional VBR and functional-structural BBR results. The three methods are: functional-functional VBR realignment with a full brain mask, functional-functional VBR realignment with a smaller brain mask and functional-structural VBR realignment. These analysis methods are discussed in detail in the Section 2.3.1.

#### 3.2.1 tSNR analysis of 7T fMRI data

The tSNR results of the three subsidiary methods are plotted alongside the two main methods in Figure 5, Panel A. The functional-structural BBR realignment (leftmost bar) is significantly better than the other four methods. The results from the functional-functional VBR realignment using SPM with the two masks (full brain and smaller brain, middle and second bar from the right) are very similar to that of the standard VBR realignment results with no mask applied (second bar from the left). Indeed, Wilcoxon signed rank test also shows no significant difference between the VBR realignment with and without mask in SPM, indicating that there is no significant benefit of removing non-brain voxels.

**Figure 5:**
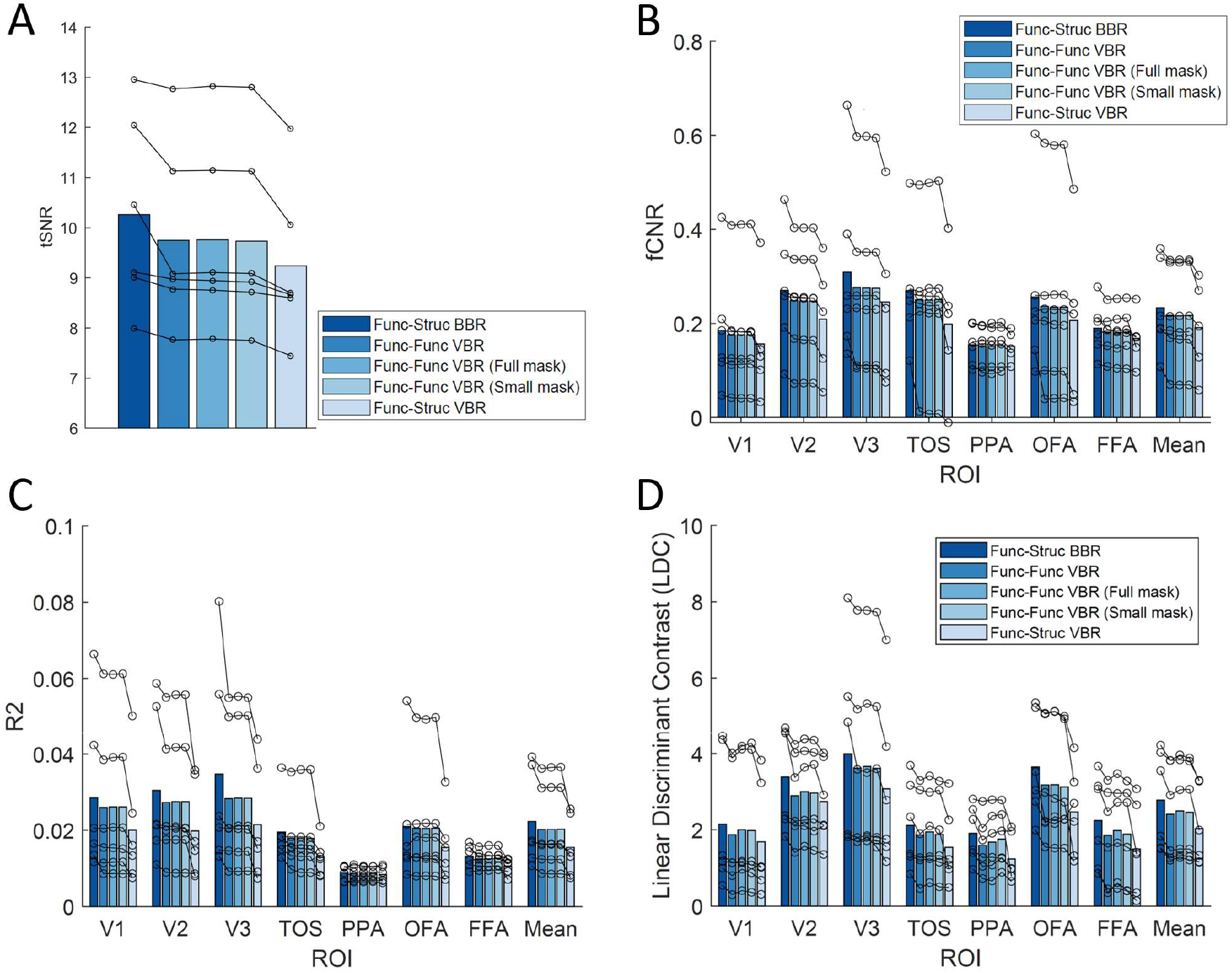
These plots compare the five different realignment methods across multiple metrics-tSNR (Panel A), fCNR (Panel B), R2 (Panel C) and Linear Discriminant Contrast (Panel D). Each set of connected circles indicate single participant results while the bar shows the group average.

When the functional-structural VBR realignment process (rightmost bar) is used, the tSNR results become significantly worse than both the functional-structural BBR realignment and the functional-functional VBR realignment results. This result showed that using VBR realignment to the structural is a worse method of realignment compared to standard realignment methods, and that the benefit of BBR realignment did not arise due to the functional-structural nature of the realignment process, but rather an inherent benefit of the BBR cost function, which emphasizes accurate registration of cortical boundaries.

#### 3.2.2 fCNR and R2 analysis of 7T fMRI data

Subsidiary analyses on the fCNR and R2 measures produced qualitatively similar results as the tSNR results (Figure 5, Panels B and C). The results using functional-functional VBR realignment with either mask (middle and second bar from the right) were similar to that of VBR realignment without mask (second bar from the left) and no significant difference was detected across all ROIs between the three methods via pairwise comparison using Wilcoxon signed-rank test for both fCNR and R2. Moreover, comparisons between the results from BBR realignment (leftmost bar) and the two functional-functional VBR realignments with masks yielded similar results to that of the comparison between BBR realignment and functional-functional VBR realignment without mask, with significant improvements in the early visual areas but no significant differences in the higher visual areas. Functional-structural VBR realignment (rightmost bar) produced results that were significantly worse than either of the main methods for most ROIs for both R2 and fCNR analysis. These results concur with the tSNR results that BBR is the best realignment method of all five used, at least for high-resolution.

#### 3.2.3 LDC analysis of 7T fMRI data

Subsidiary analysis using the multivariate LDC measure shows that utilizing a mask (middle and second bar from the right) improves the quality of the data relative to VBR realignment without mask (second bar from the left) slightly. This improvement is consistent, but not significant, across all ROIs except OFA and does not fully account for the differences between the main BBR and VBR realignment methods. Functional-structural VBR realignment (rightmost bar) produced results that are significantly worse than all other methods for all ROIs. The full results were plotted in Figure 5, Panel D.

## 4 Discussion

Retrospective motion correction (RMC) is a critical step for ensuring data quality. Our results show that BBR realignment outperforms more conventional VBR methods for realigning the grey-matter portion of 7T submillimetre data. With an increasing focus in functional activations across different cortical layers and fine-scale functional specialization, it is important to ensure proper data realignment to prevent the masking of real effects or being misled by false positives.

Initial comparisons of 7T submillimetre data using the two main methods (BBR realignment and standard whole-image VBR) showed a benefit of using BBR realignment and this benefit was observed across all four metrics used, tSNR, fCNR, R2 and LDC. All benefits were shown to be significant according to Wilcoxon signed-rank testing when averaged across all ROIs. Probing individual ROIs with fCNR and R2 showed greatest numerical improvements in ROIs near the surface of the brain, namely the early visual areas. This agrees with the tSNR comparison heatmap, which showed the greatest benefit of BBR realignment being on and near the surface of the brain. Restricting our analysis to the central region of the brain showed that both methods yielded similar tSNR results. This is expected since BBR is driven by realigning the boundaries of the brain and hence the largest benefit should be observed on and near the boundaries. However, in the LDC analysis, all ROIs showed significant improvements when BBR realignment is utilized. Given that LDC, which combines information across multiple voxels, is likely be the most sensitive metric, we can interpret these results to mean that while the major benefits of BBR realignment were localized to the brain’s surface, there were also more subtle improvements in other brain regions.

Repetition of the tSNR analysis on 3T data showed no difference between realignment methods for 1.5mm isotropic data, plus a significant decrease in tSNR for BBR realignment for 3mm isotropic data. This is in line with our expectation that BBR realignment should be most beneficial at high resolutions. Since BBR uses the brain boundaries to drive realignment, the benefit would be most apparent at higher resolutions, where the boundaries are more clearly defined. The lack of fine detail in 3mm isotropic voxels would mean that the BBR algorithm would not be able to accurately identify boundaries in the fMRI data, thus potentially leading to misalignments. Moreover, the smaller voxels at higher resolution would also cause the data to be more sensitive to small differences in realignment. Geometric distortions due to field inhomogeneities are worse at higher field strengths and this could explain why BBR is more beneficial at 7T relative to 3T.

We then attempted to probe for the source of the benefit for BBR realignment. By running standard SPM realignment with a full brain mask and also with a smaller brain mask, we obtained similar results to standard realignment without a mask for univariate analysis. LDC analysis on functional-functional VBR realignment with masks showed a slight benefit of masking over simple functional-functional VBR but was still significantly worse than functional-structural BBR realignment. Taken together, these results show that masking out non-brain voxels achieves a slight benefit on realignment, but only when using a more sensitive multivariate measure. This benefit is small though and not sufficient to explain the much larger overall advantage of BBR realignment, suggesting that the advantage of BBR realignment does not simply reflect the smaller subset of brain voxels used but rather reflects an inherent improvement due to BBR cost function used.

Functional-structural VBR generated much poorer realignment of data as compared to the other four methods. This is reflected by a significant decrease in tSNR, as well as significantly worse fCNR and R2 values across most ROIs. This is in line with our expectations, since SPM coregistration function is not designed for the purposes of time series realignment. Nonetheless, it confirms that the advantage of BBR realignment is inherent to the usage of BBR cost function, and not an artefact arising from realigning to the structural rather than between fMRI volumes.

Given that functional-structural VBR was the worst performing realignment method, it is worth considering if we could attempt functional-functional BBR. This would allow us to utilize the benefit of BBR cost function, while potentially removing the cost of a functional-structural coregistration across modalities (e.g, from images that may have different spatial distortions). However, BBR requires at least one of the images to have good definition of grey matter boundaries (normally the higher-resolution structural image) and we believe that the fMRI volumes do not typically have sufficient contrast to define those boundaries for matching. Moreover, in the Freesurfer implementation of BBR, we need to generate a surface reconstruction for the definition of boundaries. This requires either a structural image from MP2RAGE or a structural-like image from newer methods such as multi-inversion-recovery time echo planar imaging (MI-EPI) (Kashyap et al., 2018).

Our study has shown that BBR realignment is beneficial for 7T submillimetre data, especially if the region of interest for the study is near the surface of the brain. We also demonstrated that the benefits of BBR realignment is inherent to that of its realignment cost function and not due to other differences from the standard realignment approach. However, there are definitely limitations to our study. Firstly, we used a relatively unconventional field of view (FOV) due to the need to capture both higher and early visual areas with minimal repetition time (TR). Future studies using different FOVs could help further establish the advantages of using BBR realignment. Secondly, BBR realignment does not, on its own, deal with other artefacts caused by head motion, such as within-volume motion and interactions with field inhomogeneities, which cause non-rigid deformations of the image. Slice-based PMC (Huang et al., 2018; Schulz et al., 2014)is necessary to handle within-volume motion, while more sophisticated methods would be needed to model field inhomogeneities (Andersson et al., 2003; Chambers et al., 2015; Yarach et al., 2015).

## 5 Conclusion

As the field shifts towards higher resolutions and smaller voxels, participant motion during fMRI will remain an important and pertinent problem. In this paper, we presented results that show BBR realignment of fMRI volumes helps to remove inter-volume motion for fMRI time sequences and thereby improves the quality of the data, as measured by four different metrics (tSNR, fCNR, R2 and LDC). We believe that this, together with other motion correction tools, will be critical as we move towards higher resolutions.

